# Satiation State-dependent Dopaminergic Control of Food Seeking in Drosophila

**DOI:** 10.1101/240697

**Authors:** Dan Landayan, David S. Feldman, Fred W. Wolf

**Affiliations:** Quantitative & Systems Biology, University of California, Merced Merced, CA 95343 USA; Molecular Cell Biology. University of California, Merced Merced, CA 95343 USA

## Abstract

Hunger evokes stereotypic behaviors that favor the discovery of nutrients. The neural pathways that coordinate internal and external cues to motivate food seeking behaviors are only partly known. Drosophila that are food deprived increase locomotor activity, are more efficient in locating a discrete source of nutrition, and are willing to overcome adversity to obtain food. Here we developed a semi-naturalistic assay and show that two distinct dopaminergic neural circuits regulate food-seeking behaviors. One group, the PAM neurons, functions in food deprived flies while the other functions in well fed flies, and both promote food seeking. These satiation state-dependent circuits converge on dopamine D1 receptor-expressing Kenyon cells of the mushroom body, where neural activity promotes food seeking behavior independent of satiation state. These findings provide evidence for active food seeking in well-fed flies that is separable from hunger-driven seeking.

## Introduction

The neural mechanisms that regulate feeding motivation are ancient, fundamental for survival, and under complex regulation, and yet they remain partially defined and understood. Feeding motivation is classically divided into pre-ingestive and consummatory phases^1,2^. In the pre-ingestive phase, nutritional deficits cause release of hormonal signals that act on the brain to bias behavioral states towards seeking food, including heightened attention to food-related environmental cues, increased locomotion, and suppression of incompatible behaviors such as sleep. Once a nutritional source is encountered, homeostatic mechanisms in concert with sensory and nutrient detectors cause a cessation of locomotion and engagement of motor programs for food intake. Both pre-ingestive and consummatory phase behaviors are motivated and goal-directed. However, the goals and the conditions for their completion are different, suggesting that the neural circuits controlling each phase are also different. Defining the neural mechanisms of feeding motivation is important in part because the dysregulation of feeding behavior is intimately tied to obesity and eating disorders, as well as to other pathological alterations of motivation, including drug addiction^3,4^.

Simpler organisms such as Drosophila hold promise for uncovering the neural circuit mechanisms for motivated feeding behavior. In Drosophila, feeding behavior studies have focused mostly on the consummatory phase, and have revealed satiation state-dependent effects on sensory^5–7^, motor^8–10^, and central processing of feeding^11–14^. Appetitive associative conditioning with feeding has defined detailed neural circuits implicated in reward learning^15–18^. Drosophila behavioral studies of the pre-ingestive phase have focused mostly on sensory perception of appetitive stimuli, including odor tracking, satiation state-dependent olfactory acuity, and search strategies^19–23^. Here we report the development of a semi-naturalistic assay for innate pre-ingestive behaviors in Drosophila, in which flies search in an open arena for a discrete source of food. Semi-naturalistic assays may offer advantages over task-specific assays in defining how complex information is processed to drive behavior. We demonstrate specific roles for distinct dopaminergic neural circuits in the well-fed and food-deprived states for regulating food seeking behavior.

## Results

### Parametric Analysis of Drosophila Food Seeking Behavior

We developed a semi-naturalistic paradigm to measure various aspects of food seeking in freely behaving flies. Flies placed into a translucent arena (**Fig. 1A**) are tracked with a video camera (**Fig. 1B**). After a set acclimation period, a small volume of food is introduced at the center of the arena. Increasing lengths of food deprivation (wet starvation with water only) increased the number of flies in contact with the food, the food occupancy rate (**Fig. 1C**). Locomotor speed in the absence of food increased with increasing lengths of food deprivation time (**Fig. 1D**). Introduction of food into the arena rapidly decreased the locomotor speed of food deprived flies that were not in contact with the food source. Food intake scaled with deprivation time, as measured in a separate assay that minimizes seeking time (**Fig. 1E**). For subsequent experiments, ‘food-deprived’ indicates 16–20 hr of a water only diet, unless otherwise noted.

**Figure 1.**
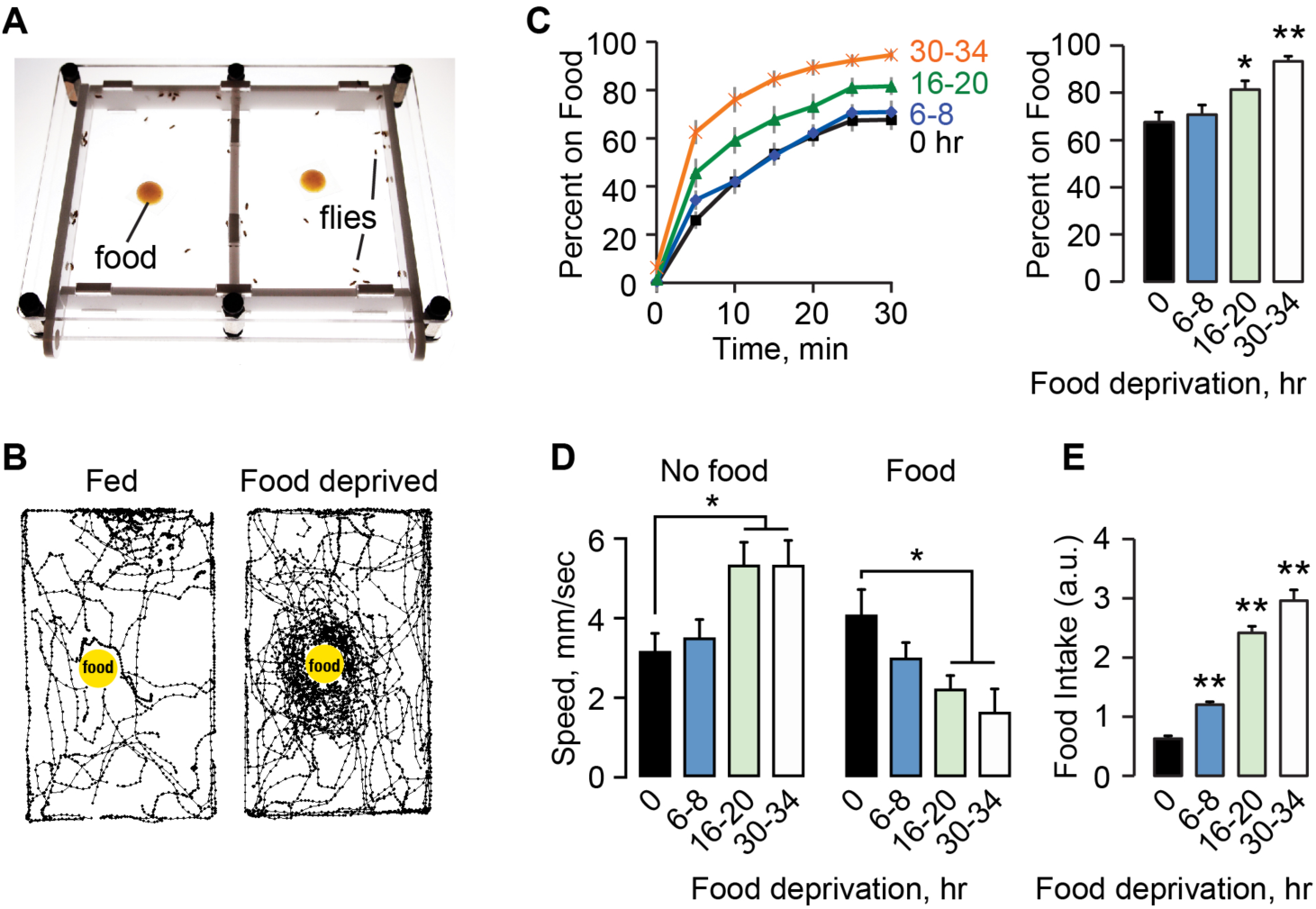
Food deprivation effect on food seeking behavior. **A**. Two-sided chamber for food seeking assays. Flies and 200 ul of cornmeal molasses food on Parafilm placed in each chamber via sliding side doors. The chamber is lit from below. Fly locomotion is recorded from above. **B**. 10 sec locomotor traces of 20 flies each filmed soon after addition of food (yellow dot). **C**. Left: The percent of flies on food over time for a food deprivation time course. Right, food occupancy averaged at 25–30 min. P<0.0001, ANOVA/Bonferroni comparison to 0 hr. n=17–18 groups. **D**. Locomotor speed. Left, speed at 20 min of acclimation, without food. Right, speed averaged over 0–10 min after food introduction. P=0.0091 no food, P=0.0066 food, ANOVA/Bonferroni compared to 0 hr. n=9–15 groups. E. Intake with increasing food deprivation time. P<0.0001, ANOVA/Bonferroni comparison to 0 hr. n=9 groups. *P<0.05, **P<0.01.

### Sensory and Nutritional Inputs to Food Seeking

We tested for the role of olfaction, taste, and vision in food seeking in food-deprived flies (**Fig. 2A**). Neither genetic nor surgical ablation of food odor-detecting neurons - olfactory coreceptor mutant *Orco^1^* or removal of the third antennal segment - affected food seeking^24,25^.

**Figure 2.**
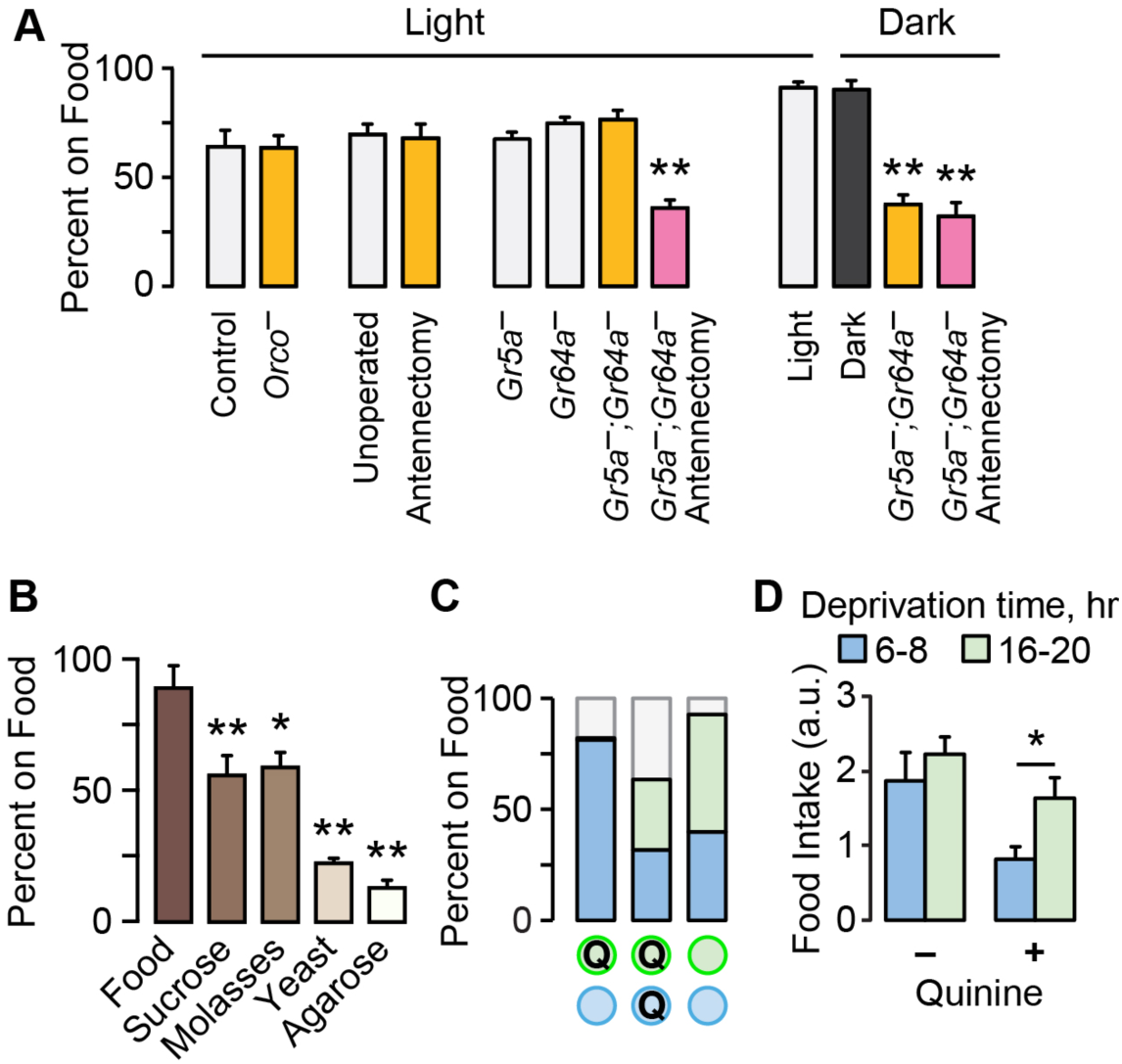
Environmental and sensory information in food seeking. **A**. Food occupancy following sensory ablations in 16–20 hr food deprived flies. Antennectomy is surgical removal of the third antennal segment. *Orco^−^* flies lack the Orco olfactory coreceptor; *Gr5a^−^* and *Gr64a^−^* are taste receptor mutants. P<0.0001 for both Light and Dark, ANOVA/Bonferroni compared to control, n=8–12 groups. **B**. Occupancy of 16–20 hr food deprived flies to agarose with the indicated food component. P<0.0001, ANOVA/Bonferroni comparison to Food. n=4–5 groups. **C**. Two-choice tests with unadulterated (open circles) and 10 mM quinine food (Q). n=5 groups. **D**. Flies consumed greater quantities of quinine food (3 mM) when food-deprived for 16–20 hr (long) versus 6–8 hr (short). P=0.0251, Mann Whitney test, n=12. *P<0.05, **P<0.01. See also Figure S1.

Similarly, flies lacking a subset of sugar sensing taste receptors had no effect on food seeking for sucrose. These experiments suggested that flies may use more than one sensory modality when seeking nearby food. Flies with both ablated antennae and taste receptor mutations showed decreased food occupancy, suggesting coordination between olfaction and taste. Food seeking remained robust in complete darkness. However, taste receptor mutant flies showed reduced food occupancy in total darkness, and additionally removing olfactory input did not further reduce occupancy. These results indicate that flies use a combination of taste, olfactory, and visual cues to find and occupy a discrete food source.

Flies may seek one or more food constituents. Food deprived flies were most attracted to complete food, then sugars, and then protein (**Fig. 2B**). When given a binary choice, flies preferred complete food over any other option, and preferred sugars over yeast (**Supplementary Fig. S1**). Similarly, flies occupied sweet and nutritious sucrose more than either sweet-only sucralose or nutritious-only sorbitol (**Supplementary Fig. S1**). Finally, nutrition appears to be important for switching the locomotor state of food deprived flies: flies slowed in the presence of sucrose or D-glucose, whereas they did not in the presence of sucralose or L-glucose (**Fig. S1D**). These findings suggest that sweetness is a mechanism that captures flies on a food source, and that nutritional content of the food source is important for fully switching flies from the pre-ingestive to consummatory phase of food seeking.

A characteristic of motivated behavior is the willingness to overcome negative consequences. Flies will eat substantially less food when it is adulterated with bitter compounds, and this scales with satiation state^13^. In a binary choice competition, food deprived flies occupied quinine-containing food, but only if there was no better choice (**Fig. 2C**). Furthermore, food intake was less suppressed by quinine with longer deprivation (**Fig. 2D**). We used a sucrose food source for all subsequent experiments.

### Role of Dopaminergic Neurons in Food Seeking

Dopaminergic neural circuits are critical for motivation, reward, and food seeking in mammals, and for many similar functions in flies^26^. To test the role of dopamine in food seeking in flies, we acutely inactivated and activated subsets of dopamine neurons in fed and food-de-prived flies and assessed occupancy of sucrose. Dopamine neurons group into several discrete anatomical and functional clusters in the adult fly brain (Fig. 3E). *TH-Gal4* labels most dopamine neuron clusters, but is largely absent from the PAM (protocerebral anterior medial) cluster of approximately 100 dopamine neurons. *0273-Gal4* labels most or all dopamine neurons in the PAM cluster but not in other dopamine neurons. Acutely blocking transmitter release in *TH-Gal4* neurons with the temperature-sensitive dynamin Shibire (Shi^ts^) had no effect on food occupancy in food deprived animals (**Fig. 3A**). Food occupancy was decreased when *TH-Gal4* neurons were transiently inactivated in fed animals. There was no effect of inactivation on locomotor activity (**Supplementary Fig. S2**). Conversely, inactivation of *0273-Gal4* neurons specifically decreased food occupancy in food deprived animals. *DAT-Gal80* (also named *R58E02-Gal80)* expresses the GAL4 inhibitor GAL80 exclusively in PAM neurons: *DAT-Gal80* blocked the *0273>Shi^ts^* food occupancy phenotype (**Fig. 3A**). Finally, chemical depletion of dopamine with 3-iodotyrosine also decreased food occupancy, indicating that dopamine is a neurotransmitter for food seeking (**Supplementary Fig. S2**). Thus, dopamine neurons in the *TH-Gal4* pattern promote food occupancy in fed animals, and PAM dopamine neurons in the *0273-Gal4* pattern promote food occupancy in food deprived animals.

**Figure 3.**
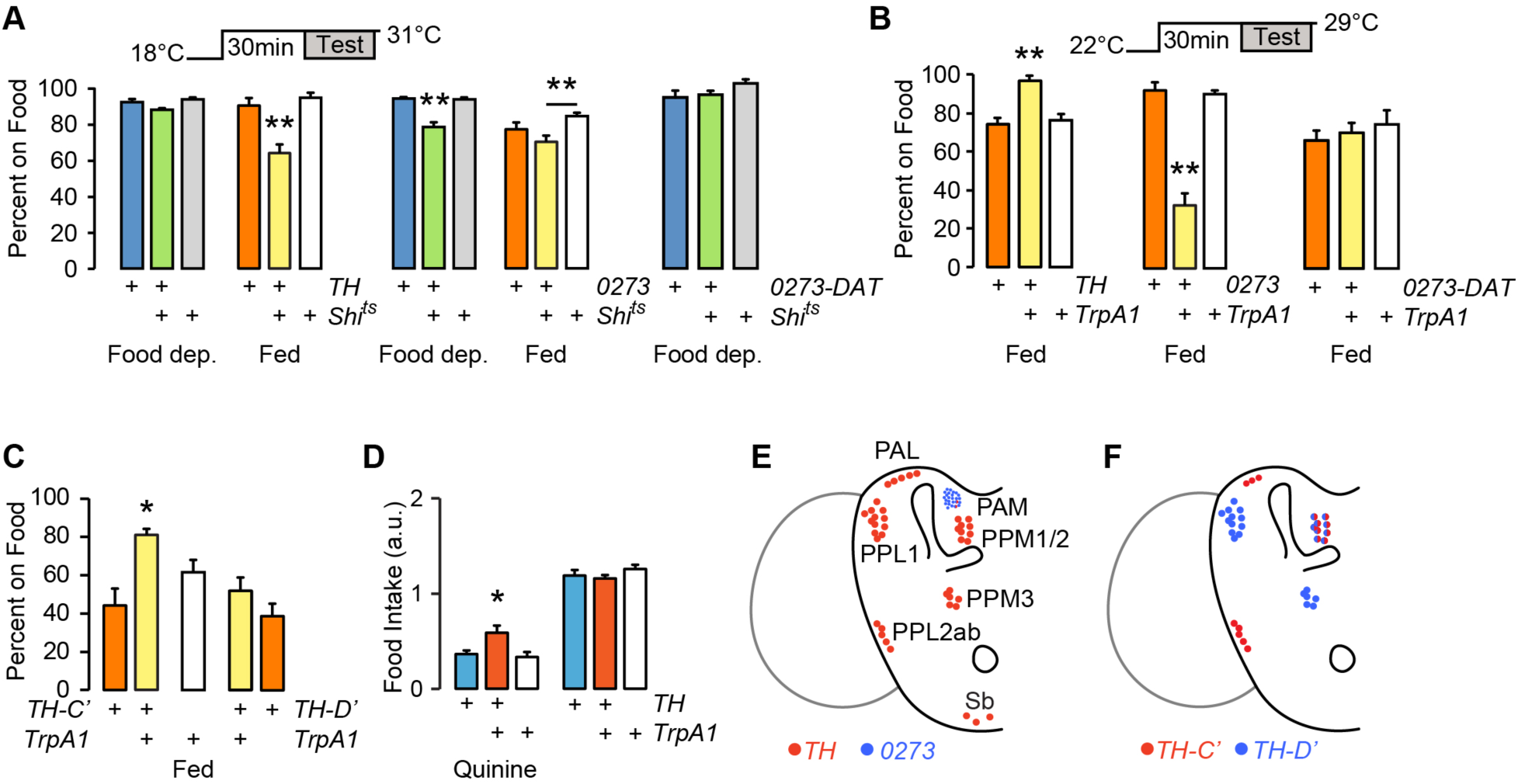
Satiation state-dependent effects of dopamine neuron activity on food seeking. **A**. Acute inactivation of dopamine neurons with Shibire^ts^ (*Shi^ts^*), food occupancy in fed and 16–20 hr food-deprived flies. P=0.0012 ANOVA/Tukey’s, n=8–11 groups with *TH-Gal4*. P=0.0001 Kruskal-Wallis/Dunn’s, n=8–10 groups food deprived; P=0.0139 ANOVA/Tukey’s, n=8–9 groups fed, with *0273-Gal4. 0273-DAT*: *0273-Gal4* with *R58E02-Gal80* to specifically block GAL4 activity in the PAM cluster dopamine neurons. n=6 groups. **B**. Acute activation of dopamine neurons in fed flies, food occupancy. P=0.0002, ANOVA/Tukey’s, n=8–11 groups with *TH-Gal4*. P=0.0002, Kruskal-Wallis/Dunn’s, n=8 groups with *0273-Gal4. 0273-DAT:* n=8 groups. **C**. Acute activation of subsets of *TH-Gal4* neurons, food occupancy in fed flies. P=0.0002, ANOVA/Tukey’s, n=8–11 groups. **D**. Food intake in 4–6 hr food-deprived flies. P=0.0053, ANOVA/Tukey’s, n=15–19 groups. **E**. Dopamine neuron clusters in the adult brain that express *TH-Gal4* and *0273-Gal4*. **F**. Dopamine neurons that express *TH-C’-Gal4* and *TH-D’-Gal4*. *P<0.05, **P<0.01. See also Figure S2.

To test if dopamine neurons are permissive or instructive, we acutely activated them using the temperature-sensitive cation channel TrpA1. Consistent with an instructive role, activating *TH-Gal4* neurons in fed flies increased food occupancy (**Fig. 3C**). Fed *0273>TrpA1* flies showed a marked decrease in food occupancy, and this was due to PAM dopaminergic activation in the *0273-Gal4* pattern. To identify the relevant neurons in the *TH-Gal4* pattern, we used transgenes that differentially label specific clusters of dopamine neurons (**Fig. 3F**)^17^. Activation of patterns that included the PPL2ab, PPM2, and PAL, but not the PPL1, PPM1, or PPM3 dopamine neuron clusters increased food occupancy in fed flies (**Fig. 3C**). To test if the identified dopaminergic neurons may regulate feeding motivation, we activated *TH-Gal4* neurons in mildly (4 hr) food-deprived flies. Under these conditions, activation of *TH-Gal4* neurons specifically increased consumption of quinine adulterated food (**Fig. 3D**). Taken together, these experiments are consistent with dual roles for dopamine in food-seeking behavior: a PAM dopamine neuron-mediated promotion of food seeking in the food-deprived state, and a *TH-Gal4* dopamine neuron-mediated promotion of food seeking in the fed state. In the fed state, PAM dopamine neurons can block food seeking.

### Dopamine Receptor Regulation of Food Seeking

*Dop1R1* encodes a D1-like dopamine receptor that functions in motivation-related behaviors, including arousal state, drug reward, and learning and memory^27–29^. We tested flies with strongly reduced expression of *Dop1R1* for food seeking behaviors. Food-deprived *Dop1R1* mutant flies were hyperactive and appeared to ignore food (**Fig. 4A**). Moreover, *Dop1R1* mutant food occupancy was reduced when fed or food deprived (**Fig. 4B**). Loss of the dopamine D2-like receptor *D2R* did not affect food occupancy, but restored normal food occupancy to *Dop1R1* mutants. These data suggest that *Dop1R1* promotes food seeking, and that an opposite role for *D2R* is uncovered in the absence of *Dop1R1*. Food intake was unaffected in food-deprived flies of these genotypes (**Supplementary Fig. S3**).

**Figure 4.**
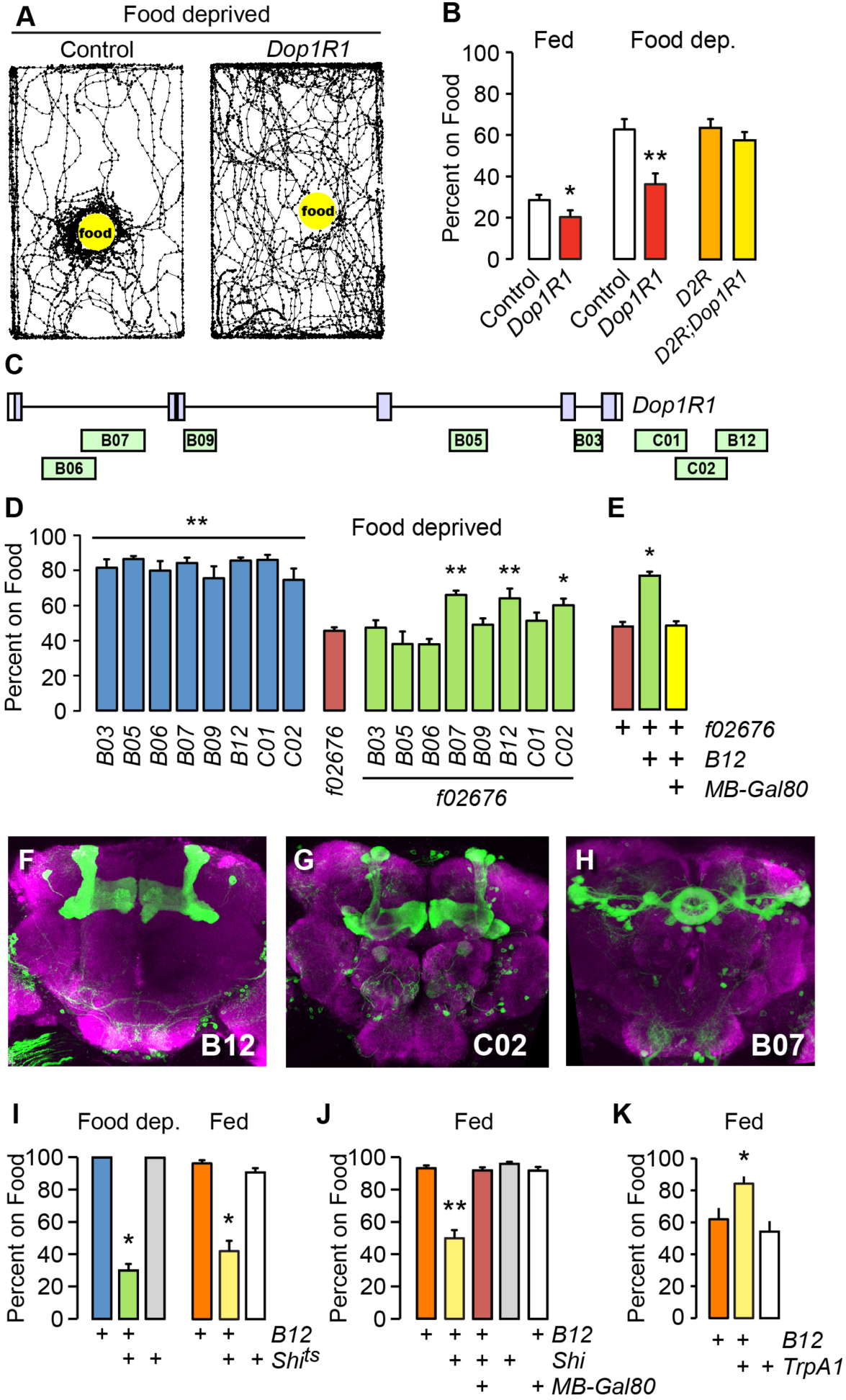
Dopamine receptor-expressing neurons in the mushroom body control food seeking. **A**. Locomotor traces of food-deprived flies 5 min after addition of food. *Dop1R1* mutant *f02676* vs. Berlin genetic background control. **B**. Food occupancy for the indicated genotypes that were fed or food deprived. t-test P=0.0492 fed (n=16–20 groups), P=0.001 food deprived (n=16–20 groups). *D2R*: the loss-of-function mutation *f06521*. **C**. Location of *Dop1R1* enhancer fragments. **D**. Genetic rescue of *Dop1R1* mutant food occupancy in 16–20 hr food deprived animals. *Dop1R1-Gal4* strains (blue) were made heterozygous in *f02676* homozygotes (rescuing configuration, green). P<0.0001 ANOVA/Bonferroni’s comparison to *f02676*, n=8–16 groups. **E**. Inclusion of *MB-Gal80*, preventing GAL4 activity in the mushroom bodies blocks *B12* rescue. P<0.0001 ANOVA/Tukey’s, n=10–19 groups. **F-H**. Expression pattern of *Dop1R1-Gal4* strains (CD8-GFP, green), and bruchpilot (magenta) to show the synaptic neuropil. **I**. Acute silencing of *B12 Dop1R1-Gal4* neurons with *Shi^ts^*, food occupancy, food deprived and fed. Food deprived: P<0.0001 Kruskal-Wallis/Dunn’s, n=4 groups. Fed: P=0.0002 Kruskal-Wallis/Dunn’s, n=7–8 groups. J. Addition of *MB-Gal80* in *B12 Dop1R1-Gal4>Shi^ts^* fed flies, food occupancy. P<0.0001 Kruskal-Wallis/Dunn’s, n=6–10 groups. K. Activation of *B12 Dop1R1-Gal4* neurons in fed flies increased food occupancy. P=0.0054, ANOVA/Tukey’s, n=7–9 groups. *P<0.05, **P<0.01. See also Figure S3.

### The Mushroom Bodies Promote Food Seeking Independent of Satiation State

We performed genetic rescue experiments to ask where *Dop1R1* functions for food seeking in food deprived flies. To bias the rescue towards functionally relevant brain regions, we utilized *Dop1R1-Gal4* strains that expressed GAL4 under the control of short non-coding genomic DNA fragments cloned from the *Dop1R1* locus (**Fig. 4C**). Food occupancy was partially rescued when *Dop1R1* was expressed with three different *Dop1R1-Gal4* strains in food-deprived *Dop1R1* mutants: *B07, B12*, and *C02* (**Fig. 4E**). Anatomical analysis of the expression patterns for the rescuing *Dop1R1-Gal4* drivers revealed expression overlap. In the *B12* and *C02* strains, the mushroom bodies were prominently labeled, as were regions of the central complex, including the fan-shaped body and protocerebral bridge (**Fig. 4F,G**). The *B07* strain prominently labeled the ellipsoid body of the central complex (**Fig. 4H**). We failed to rescue *Dop1R1* mutant food occupancy using GAL4 drivers that label the ellipsoid body, fan-shaped body, or the protocerebral bridge (not shown). By contrast, decreasing GAL4 activity with mushroom body-specific expression of GAL80 eliminated *B12* rescue of the *Dop1R1* mutant food occupancy phenotypes (**Fig. 4E**). Moreover, restoring *Dop1R1* with the mushroom body-specific driver *MB247-Gal4* rescued *Dop1R1* food seeking (**Supplementary Fig. S3**). Thus, *Dop1R1* expression in the mushroom bodies is sufficient to promote food seeking in food deprived animals.

We next tested the role of neurotransmission in Dop1R1-expressing mushroom body neurons in food seeking. Similar to loss of *Dop1R1*, acute blockade of synaptic output in *B12* neurons decreased food occupancy in both fed and food-deprived flies (**Fig. 4I**). Importantly, this effect also localized to the mushroom bodies (**Fig. 4J**). *B12>Shi^ts^* flies also showed reduced locomotion, however this phenotype persisted when the mushroom body neurons were subtracted from *B12* (**Supplementary Fig. S3**), suggesting that distinct Dop1R1 neurons control food occupancy and locomotion. Finally, acute activation of *B12* neurons in fed flies increased food occupancy (**Fig. 4K**). Taken together, these results indicate that the activity of Dop1R1-expressing mushroom body neurons promote motivated food seeking in both the fed and food-deprived state.

## Discussion

Distinct dopaminergic circuitry promotes food seeking under well fed and food deprived conditions. Dopamine neurons in the TH-C’ pattern promote seeking in well fed flies, and dopamine neurons in the PAM cluster promote seeking in food deprived flies. The PAM neurons likely function directly upstream of Dop1R1-expressing neurons of the mushroom body that promote food seeking in both the fed and food-deprived states. These circuits function in food seeking under semi-naturalistic conditions, where flies can freely perform many steps of food seeking behavior. Understanding how these dopaminergic circuits contribute to discrete steps of feeding behavior, from local search through to repletion and disengagement from a food source, will help define how motivational states transition from task to task.

### Roles of Dopamine in Appetitive Behaviors

Dopaminergic neurons are critical for many appetitive and aversive behavioral responses across animal species. Dopamine may act as a salience, arousal, or attention signal that gives importance to specific valence information arriving from other circuit elements^26,30,31^. In rodents, genetic, pharmacological, and lesioning studies indicate that striatal dopaminergic pathways can selectively function in the pre-ingestive phase to promote food seeking^30,32,33^. We found that acute activation of dopamine neurons in fed flies increased food occupancy, yet it did not cause increased food intake. Likewise, genetic elimination of the Dop1R1 receptor decreased food occupancy without affecting food intake. In contrast, inactivation of Dop1R1 receptor neurons decreased food intake in the food-deprived state, possibly reflecting their key role in integrating sensory and internal state information. These findings suggest that dopaminergic pathways promote pre-ingestive food seeking. However, the role of dopamine is more complex. For example, the PAM dopamine neurons are activated by ingestion of sugar, and their activation is greater in food-deprived flies, indicating that dopaminergic neurons are engaged during the consummatory phase of feeding, and they may be sensitized to responding to input during the pre-ingestive phase^17^.

Prior studies assigned dopamine to particular aspects of feeding behavior and also to motor functions that are critical to feeding^14,23^. In particular, dopamine neurons in the *TH-Gal4* pattern are implicated in controlling motor output: *TH-Gal4* neuron hyperpolarization, blocking synaptic input, interferes with motor performance and aspects of food seeking behavior in food deprived flies^23,34^. We did not detect differences in unstimulated motor activity or in the magnitude of an olfactory-stimulated startle response when we blocked synaptic output from *TH-Gal4* neurons, indicating that flies exhibited grossly normal motor behavior in our assay^35^. The differences in observed phenotypes may reflect the multifunctional roles of *TH-Gal4* dopamine neurons that are revealed by specific types of manipulation.

Which dopamine neurons are responsible for food seeking? In well-fed flies, neurons in the *TH-C’* pattern promote seeking. This pattern includes dopamine neurons in the PAL, PPM2, and PPL2 clusters, and this group of neurons was previously shown to promote female egg-laying preference on sucrose^17,36^. The neurons in these clusters project to many specific regions of the brain, and their individual functions remain largely unknown. One exception is that individual neurons in the PPM2 cluster, the DA-WED neurons, support protein consumption preference in protein deprived flies^14^. The DA-WED neurons synapse to Dop1R1 neurons in the *B03* pattern, which did not support rescue of food seeking in our experiments. These findings argue that there are distinct dopaminergic circuits in the *TH-C’* pattern that control different forms of nutrient seeking. The PAM neurons are also heterogeneous, sending projections that tile to well-defined regions of the mushroom body and to regions of the protocerebrum. Specific subsets of PAM neurons that are included in the *0273-Gal4* pattern have been implicated in various forms of appetitive learning and memory, however their inactivation did not impact food seeking in food deprived flies (not shown)^15,17,37–39^. This suggests that there may be further segregation of PAM dopamine neuron function, possibly according to innate and learned appetitive responses.

### Sensory Tuning of Food Seeking Motivation

Appetitive olfactory cues such as those emitted from palatable food elicit approach and can activate neurons important for feeding. Olfactory receptor neurons that respond to appetitive odors increase sensitivity through the actions of the neuropeptides sNPF and SIFamide ^21,40^. Further, neurons that release the neuropeptide NPF are activated to a greater extent in response to food odors in food-deprived flies; their activation promotes and inactivation inhibits odor attraction^41^. In well-fed larvae, the attractive odor pentyl acetate increases food intake through the actions of NPF and dopamine^11^. Therefore, food-related odors not only elicit approach behavior in a satiation state dependent manner, but also increase the activity of neurons expressing neuropeptides that regulate feeding behavior. Our results indicate that, under semi-naturalistic conditions, olfaction is important but apparently not crucial for food seeking in food-deprived flies: neither surgical nor genetic ablation of olfaction decreased food occupancy, and its role was only revealed by simultaneous partial ablation of taste responses. Further, flies were efficient in seeking odorless sucrose. Taken together, olfaction, hygrosensation, visual cues, and taste responses likely act in concert with internal cues to set the intensity of food seeking when freely behaving flies are in close proximity to a food source.

## Methods

### Strains and Culturing

All strains were outcrossed for five generations to the Berlin genetic background prior to behavioral testing. Flies were raised on standard food containing agar (1.2% w/v), cornmeal (6.75% w/v), molasses (9% v/v), and yeast (1.7% w/v) at 25°C and 70% humidity unless otherwise indicated. *Dop1R1-Gal4* (*R72B03, R72B05, R72B06, R72B07, R72B09, R72B12, R72C01*, *R72C02*) strains were generated by the FlyLight project^39^, *TH-C’-Gal4* and *TH-D’-Gal4* were from Mark Wu, *Gr5a^EP-5^* and *Gr64^a1^* were from Anupama Dahanukar, *0273-Gal4* from Daryl Gohl and Thomas Clandinin, *MB-Gal80* from Hiro-mu Tanimoto, *Orco^1^* from Leslie Vosshall, and others from the Bloomington Stock Center.

### Behavioral Measurements

Groups of 21 males were collected 1–2 days prior to the experiment. For food deprivation flies were placed into empty culture vials containing water saturated Whatman filter paper. For 3-iodotyrosine treatment flies were cultured for 30 hr with 5% sucrose/2% yeast/10 mg/mL 3-iodotyrosine (3IY), and treated an additional 16 hr with 3IY in water for food deprivation. Standard fly food was used for all experiments except where indicated. Thin-walled Plexiglas behavioral chambers were designed with two side-by-side arenas, each arena measuring 45x75x10 mm, or 85×135×10 mm for experiments with *Shibire^ts^*. Chambers were designed and built by IO Rodeo (Pasadena, CA). Flies were filmed from above at 10 fps with the arena placed on white light LED panel (Edmund Optics). Filmed flies were tracked with customized software^40^. For food occupancy, the number of flies off food was subtracted from the total number of flies and divided by total number of flies. Percent on food was calculated as the average of the last two measured time points. Locomotor activity was the average speed of all flies in 20 sec bins.

To measure food intake, 5 ml standard fly food with 2% erioglaucine (Sigma) with or without 3mM quinine was striped onto 1/4 of the inner surface of a wide fly vial, and condensation removed. 30–50 flies were introduced and the vial laid on its side so that the food edge was at the apex. After 30 min, the flies were homogenized in a volume adjusted to the number of flies and consumption was determined spectrophotometrically.

Statistical measurements were made with Prism 6.0 (GraphPad). Error bars are the SEM. Data is available upon request.

### Immunohistochemistry

Adult fly brains were fixed and immunostained as described previously^32^. Antibodies were rabbit anti-GFP (1:1000, Life Technologies), rabbit anti-Dop1R1 1:1250^32^, and nc82 (1:25, Developmental Studies Hybridoma Bank, Iowa).

## Author Contributions

DL, DSF, and FWW conceived of and carried out the experiments, and analyzed the results. FWW wrote the paper.

## Acknowledgements

We thank the members of the laboratories of Fred Wolf and Michael Cleary for advice, and Daryl Gohl and Thomas Clandinin for unpublished strains. This work was supported by grants from the NIH (AA018799), The Hellman Fellowship Fund, and the University of California, Merced. The authors declare no competing financial interests.

## Supplementary Information

### Supplementary Figures

**Figure S1, related to Figure 2.**
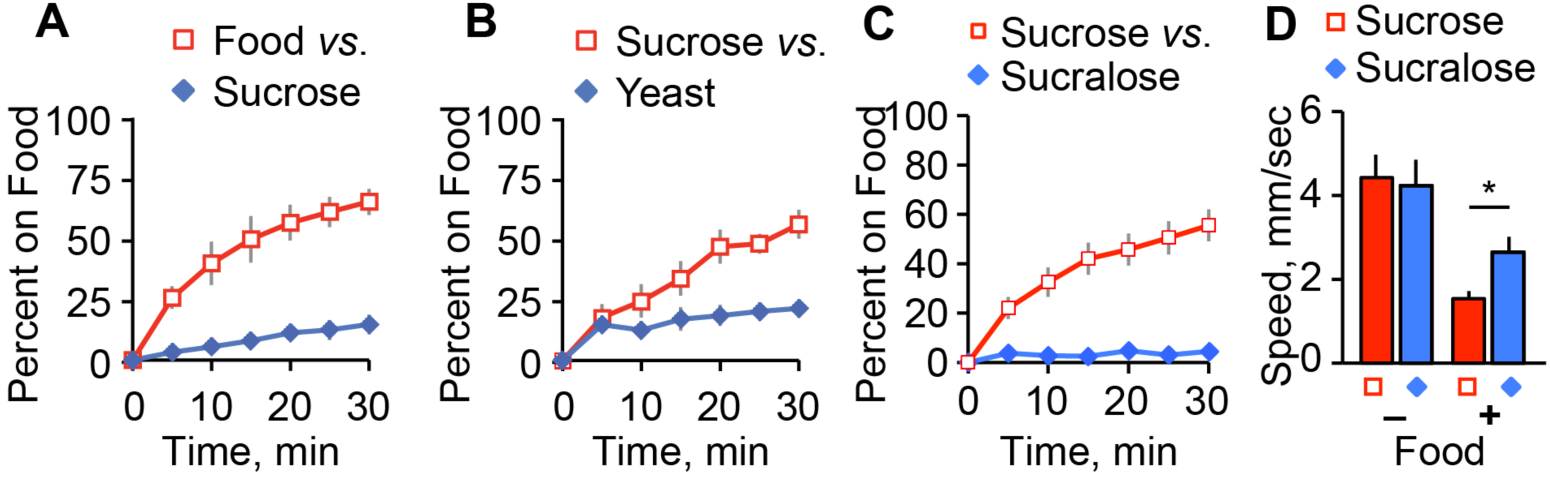
Food occupancy with binary choice. Percent of flies occupying **A**. food vs. sucrose, **B**. sucrose vs. yeast, and **C**. sucrose vs. sucralose. **D**. Locomotor speed of flies before and after addition of the indicated food source. t-test, *P<0.05. n=6–10 groups.

**Figure S2, related to Figure 3.**
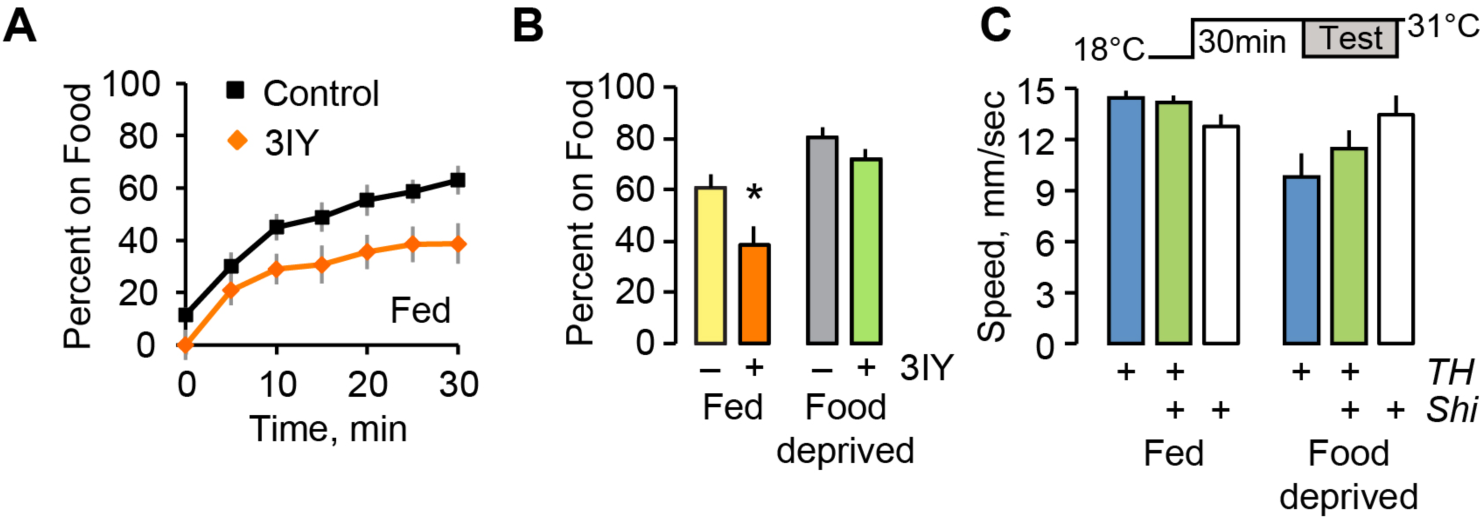
Food occupancy after overnight treatment with 10 mg/mL 3-iodotyrosine (3IY). **A**. Time course. **B**. Percent occupancy. t-test, *P<0.05. n=8 groups. **C**. Locomotor speed with acute inactivation of *TH-Gal4* neurons. n=8–11 groups.

**Figure S3, related to Figure 4.**
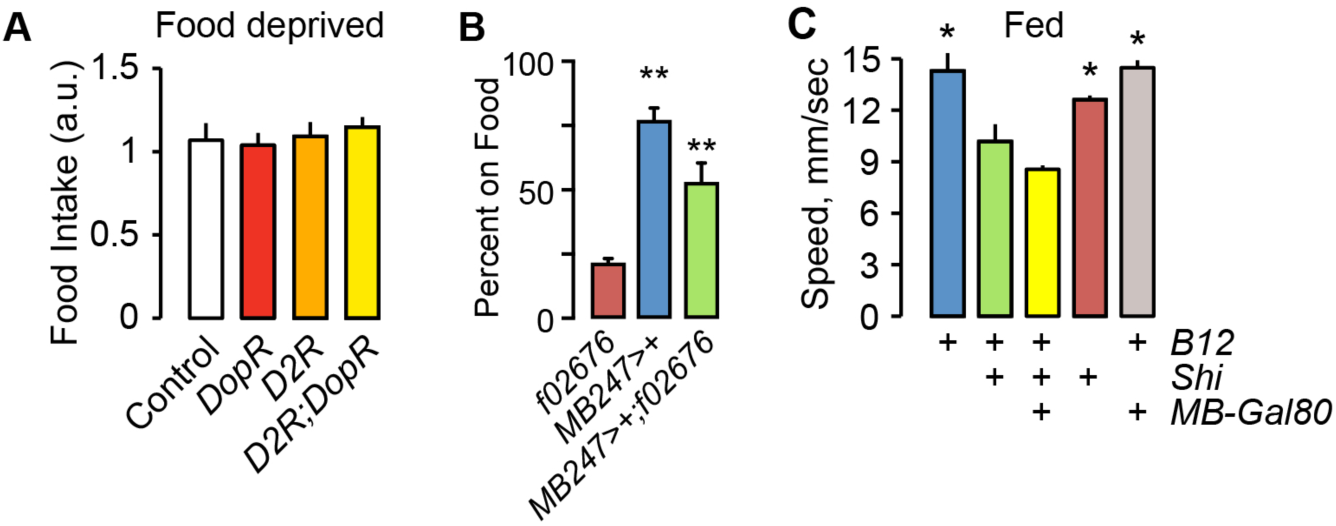
Dopamine receptor neuron manipulation. **A**. Food intake for the indicated genotypes. n=10 groups. **B**. Food occupancy for genetic rescue restricted to the mushroom bodies. P<0001, One-way ANOVA/Tukey’s. n=12 groups. **C**. Locomotor speed in fed flies of the indicated genotypes. P<0.0001 One-way ANOVA/Bonferroni compared to *B12-Gal4>UAS-Shi^ts^*. n=8–12 groups.

